# Adversarial Sequence Mutations in AlphaFold and ESMFold Reveal Nonphysical Structural Invariance, Confidence Failures, and Concerns for Protein Design

**DOI:** 10.64898/2026.02.25.708002

**Authors:** Jonathan Feldman, Maximilian Brogi, Jeffrey Skolnick

## Abstract

AlphaFold has transformed structural biology and spawned an ecosystem of derivative tools for protein design, binding prediction, and drug discovery. However, whether AlphaFold has learned generalizable biophysical principles versus template-based pattern matching remains unclear—a distinction critical for applications beyond its training context. Here, we perform a systematic adversarial evaluation of AlphaFold 3 using point and deletion mutations across 200 proteins. Remarkably, predicted structures remain invariant to mutations of up to 40% of residues—including deliberately destabilizing substitutions—and to deletions of 10%. Notably, this invariance holds even for experimentally validated fold-switching proteins that are known to adopt alternative conformations in response to such mutations, despite the fact that these proteins are small and monomeric—precisely the category where AlphaFold is expected to perform best. Confidence metrics prove unreliable, as they select the most accurate structure at most 35% of the time and correlate with the structural quality of the best available training set template. This suggests that AlphaFold’s uncertainty estimates reflect template availability more than biophysical reasoning. ESMFold exhibits greater, though still imperfect, mutational sensitivity, suggesting superior sequence-structure coupling. These findings indicate that AlphaFold may rely heavily on memorized templates rather than biophysical reasoning, with profound implications for the reliability of AlphaFold-based protein design, drug discovery, and modeling workflows.

## 1 Introduction

The release of AlphaFold 2 in 2021 fundamentally transformed structural biology [1–4]. By achieving near-experimental accuracy on a substantial fraction of protein targets, AlphaFold 2 demonstrated that deep learning could solve what had been considered one of biology’s grand challenges: predicting a three-dimensional protein structure from its amino acid sequence alone [1, 5, 6]. This breakthrough catalyzed an explosion of research across computational biology, accelerating drug discovery, enabling structure-guided protein engineering, and providing structural insights into previously intractable systems [7–10].

AlphaFold 3, released in 2024, extended these capabilities further [11]. Beyond proteins, it can model nucleic acids, small molecules, and ligands, enabling prediction of complete biomolecular assemblies [11, 12]. Its diffusion-based architecture and expanded training set yield improved accuracy, particularly for protein-protein interfaces and multimeric complexes [8, 11, 13]. As a result, AlphaFold 3 has rapidly become the de facto standard in computational structural biology, underpinning an expanding ecosystem of derivative tools and methods.

This ecosystem now extends far beyond structure prediction. Modern protein design frameworks such as RFdiffusion and ProteinMPNN rely on architectural principles and training strategies pioneered by AlphaFold [14–16]. Generative models for protein binder design—including BoltzGen, BindCraft, and variants of AlphaFold itself—use AlphaFold’s structure prediction modules as differentiable components through which gradients are backpropagated to optimize sequences [17–19]. Protein language models, which learn sequence representations without explicit structural supervision, are increasingly coupled with AlphaFold’s predictions to enable tasks ranging from fitness prediction to genomic design [20–22]. Tools for predicting binding affinity, stability changes upon mutation, and functional annotations increasingly incorporate AlphaFold-derived features or use AlphaFold as a preprocessing step [23–25]. In experimental laboratories, AlphaFold predictions now routinely guide construct design, inform crystallization strategies, and aid in molecular replacement for structure determination [26, 27].

The pervasive integration of AlphaFold into research workflows makes understanding its underlying capabilities—and limitations—a matter of broad scientific importance. If AlphaFold has learned the true biophysical principles that relate sequence to structure, then the tools built upon it inherit a robust foundation. However, if AlphaFold’s predictions are primarily driven by template matching to its library of training proteins—similar to a threading algorithm [28]—then derivative methods may be inheriting the same constraints and are effectively glued to the training structures regardless of what conformation the sequence of interest will actually adopt. The distinction is not trivial: models that lack biophysical grounding are unlikely to generalize to novel protein families, engineered sequences far from natural distributions, or sequences with substantial mutations relative to characterized homologs [8, 29–32]. There is mounting evidence that current structure prediction models struggle with such out-of-distribution inputs. AlphaFold’s performance degrades substantially on proteins with low sequence similarity to its training set [33, 34]. Its predictions for alternative conformations and fold-switching proteins often fail to reflect known structural diversity, instead converging to single dominant folds [35, 36]. When used for protein design, AlphaFold-based pipelines can generate sequences that appear well-folded in silico but fail to express, fold correctly, or exhibit intended functions in vitro [17, 37, 38]. These observations raise critical questions: To what extent has AlphaFold learned the biophysical rules governing protein folding? Does it reason about how specific residues contribute to stability, interface formation, and structural integrity? Or does it somehow recognize patterns from its training data and interpolate among known structures to essentially recapitulate the training structure?

Addressing these questions requires adversarial evaluation beyond standard held-out benchmarks, which measure predictive accuracy but do not test whether models capture causal sequence–structure relationships. A model that has learned biophysical principles correctly should respond to mutations in proportion to their structural impact—–for example, deleterious core mutations should be more disruptive than surface substitutions, and deletions more disruptive than conservative point changes—whereas a template-driven model may remain invariant despite substantial, and nonphysical, sequence divergence.

Here, we perform a systematic adversarial evaluation of AlphaFold 3 using complementary point-mutation and deletion-mutation regimes. We analyzed over 200 proteins spanning multiple categories, including proteins with and without training-set homologs and experimentally validated fold-switching proteins. We then compare AlphaFold 3 with ESMFold, a protein language–model–based method that does not rely on multiple sequence alignments (MSAs) [39], and finally, evaluate AlphaFold 3 confidence metrics in relation to those of AlphaFold 2. Together, these analyses probe AlphaFold 3’s mutational sensitivity and help distinguish learned biophysical generalization from structural memorization, with implications for the reliability of AlphaFold-based studies.

## 2 Results

### 2.1 Adversarial Point Mutation Analysis

To probe how AlphaFold 3 incorporates biophysical constraints and reality into its predictions, we conducted an adversarial point mutational analysis. A total of 200 protein sequences spanning four categories—monomer-novel, monomer-similar, multimer-novel, and multimer-similar—were provided to AlphaFold 3 for structure prediction. Here, monomeric and multimeric denote the number of chains, while “similar” indicates greater than 30% sequence homology to at least one protein in the AlphaFold 3 training set, and “novel” indicates the absence of such homology. Each sequence was then mutated at 5, 10, 20, 40, and 70% point-mutation thresholds, with structure predictions generated for every mutated variant. Mutations were intentionally selected to be maximally disruptive—for example, substituting small hydrophobic residues with large polar residues—and were biased toward positions near the center of the sequence, a region more likely to map to the protein core [40, 41]. See Methods for more details.

For each mutation level, predicted structures were compared to the original predicted structure, which served as the reference. Comparisons used both a global structural similarity metric, TM-score [42, 43], and an interfacial accuracy metric, DockQ [43, 44].

As shown in Figure 1, AlphaFold 3 maintains global fold similarity up to remarkably high levels of perturbation. Figure 1H presents a survival curve representing the fraction of structures that retain the same global fold—defined as TM-score *≥*0.5— across mutation thresholds. At 40% mutation, monomeric proteins still maintain the same global fold on average, while multimeric proteins, though they degrade in quality more rapidly, still preserve global fold correctness in over 40% of cases. This pattern holds consistently for both novel and previously seen sequences, with no discernible difference observed between proteins novel to the AlphaFold 3 training set and those with homologs in it.

**Fig. 1.**
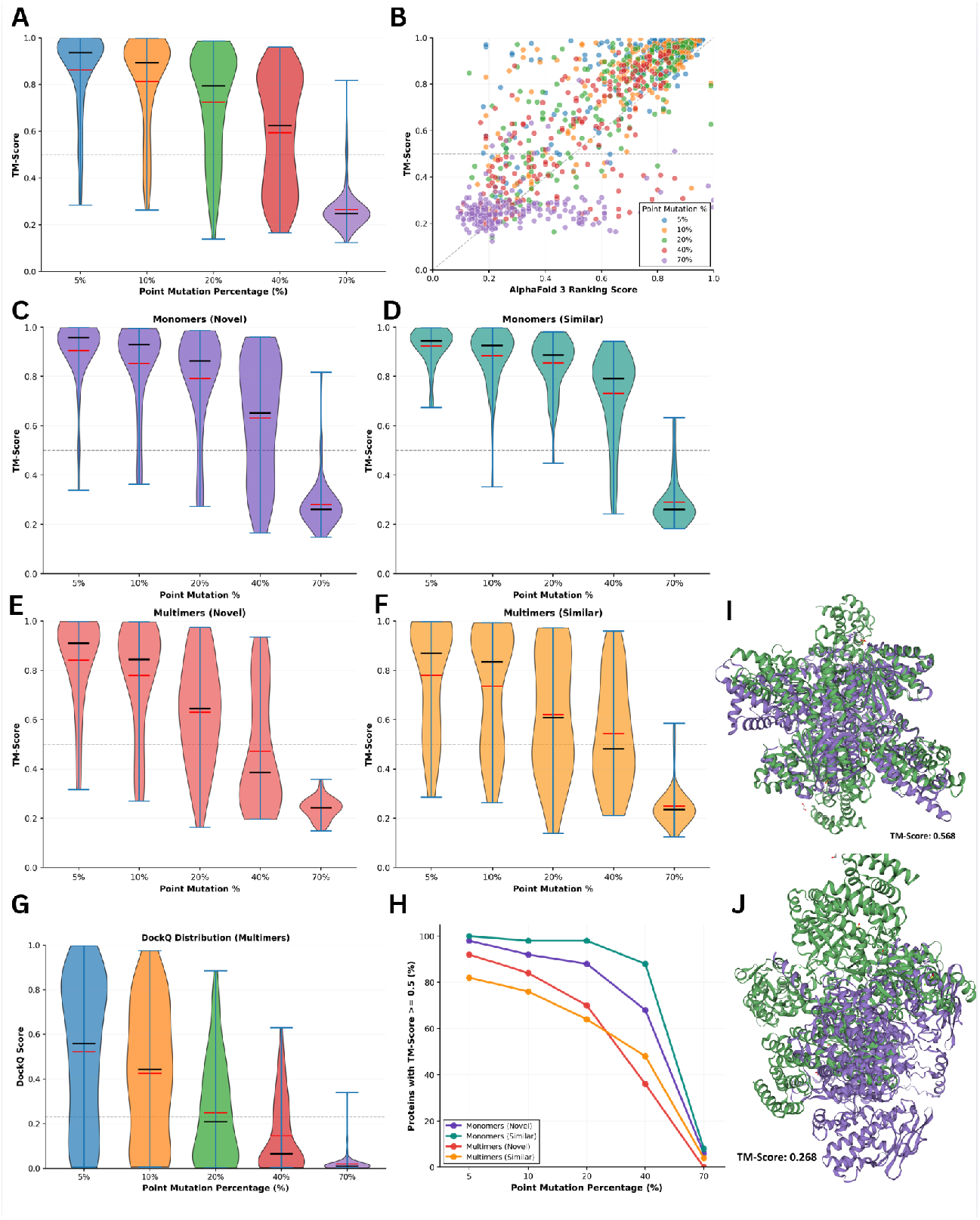
AlphaFold 3 exhibits structural invariance under adversarial point mutations. **A** Violin plots showing TM-score distributions for the full 200-protein dataset at each mutation threshold (5%, 10%, 20%, 40%, 70%), comparing mutated predictions to unmutated AlphaFold 3 predictions. **B** Correlation between AlphaFold 3 ranking confidence scores and structural accuracy (TM-score). Each point represents a single protein, colored by mutation percentage. **C–F** TM-score distributions stratified by protein category: (**C**) monomer-novel, (**D**) monomer-similar, (**E**) multimer-novel, and (**F**) multimer-similar. **G** DockQ score distributions for multimeric proteins (novel and similar bins combined), showing interfacial accuracy degradation across mutation thresholds. **H** Survival curve indicating the fraction of proteins maintaining accurate global fold (TM-score *≥* 0.5) at each mutation level. **I** Structural superposition of 7OCN with 40% mutation (purple) onto the unmutated prediction (green), showing preserved fold architecture (TM-score = 0.568). **J** Structural superposition of 7OCN with 70% mutation (purple) onto the unmutated prediction (green), showing fold destruction (TM-score = 0.268). All TM-scores and DockQ values reflect comparison to the unmutated AlphaFold 3 prediction rather than experimental structures.

The violin plots in Figure 1A and G reveal how TM-score and DockQ distributions shift across mutation thresholds. TM-scores remain remarkably high through 70% mutation, declining only gradually as sequence identity erodes. In stark contrast, interfacial geometry degrades rapidly. DockQ scores show that by 20% mutation, approximately half of the interfacial structures are incorrect with respect to the structure predicted from the original unmutated sequence, highlighting that interfacial contacts are far more vulnerable to mutational perturbation than overall fold architecture.

AlphaFold 3’s internal confidence metrics show only moderate sensitivity to these perturbations. The relationship between mutation percentage and ranking score is modest, with a Pearson correlation coefficient of *−* 0.558 and a Spearman rank correlation of *−* 0.544. Consistent with this trend, the mean AlphaFold 3 ranking score decreases gradually from 0.73 at 5% mutation to 0.62 at 20%, and falls below the low-confidence regime only at the 40% mutation level. Notably, the commonly used threshold for high-confidence predictions is a ranking score of 0.6 [4, 25]; thus, even at 20% mutation—–where a substantial portion of the protein has been deliberately altered in a highly deleterious manner—the model remains confident in its predictions on average.

We extended the point mutation analysis to fifteen experimentally validated, monomeric fold-switching proteins compiled by a previous study [45]. For each protein, mutations were directed at residues or regions that prior work has empirically shown to induce fold-switching (details on the mutation regime for each protein are provided in Methods and in the Supplementary Information). Table 1 summarizes the results. Consistent with the larger dataset, predicted structures remained highly invariant up to the 40% mutation threshold, maintaining an average TM-score of at least 0.63 relative to the original predicted structure. Even for proteins where the empirically validated fold-switching residues or regions were purposefully mutated, AlphaFold 3 exhibited little detectable structural divergence, mirroring the behavior observed in non–fold-switching proteins. This result reinforces the conclusion that AlphaFold 3 frequently fails to capture biologically plausible mutational responses, even when multiple conformations are known to exist. The AlphaFold 3 ranking score similarly remains high up to the 40% threshold, further underscoring confidence metric invariance. Hence, even for the small proteins where fold-switching occurs, AlphaFold 3 does not reliably respond to known fold-switching mutations, suggesting it should be used with caution in protein sequence optimization.

**Table 1.** Structural and confidence metrics for fold-switching proteins under point mutations. TM-score and AlphaFold 3 ranking score correlations with mutation percentage, along with mean metric values at each mutation threshold. Correlation coefficients quantify the relationship between mutation level and metric degradation. Mean values are computed across fifteen experimentally validated fold-switching proteins from Porter et al. [45].

| Metric | Correlation | Coefficient | 5% | 10% | 20% | 40% | 70% | Mean |
| --- | --- | --- | --- | --- | --- | --- | --- | --- |
| TM-score | Pearson | −0.66 | 0.77 | 0.70 | 0.63 | 0.49 | 0.31 | 0.58 |
|  | Spearman | −0.66 | — | — | — | — | — | — |
| Ranking score | Pearson | −0.57 | 0.77 | 0.74 | 0.70 | 0.62 | 0.51 | 0.67 |
|  | Spearman | −0.55 | — | — | — | — | — | — |

### 2.2 Adversarial Deletion Mutation Analysis

To further assess AlphaFold 3’s susceptibility to memorization or structural invariance, we conducted an analogous adversarial experiment using residue deletions, which are typically far more disruptive than point mutations [46]. Even a small number of deletions can cause severe destabilization or collapse of tertiary or quaternary structure [46, 47]. Accordingly, lower deletion thresholds of 1, 3, 5, and 10% were employed. As in the point-mutation experiment, deletions were biased toward centrally located residues to maximize structural disruption.

Despite the severity of this perturbation type, AlphaFold 3 again maintains global fold integrity, as shown in Figure 2. The survival plot in Figure 2H captures this clearly: even at 10% deletions, an overwhelming majority of monomeric proteins retain accurate predicted structures. Multimeric proteins decrease in accuracy more rapidly than monomers, consistent with the point mutation results, but no significant difference was observed between novel and similar proteins within either category.

**Fig. 2.**
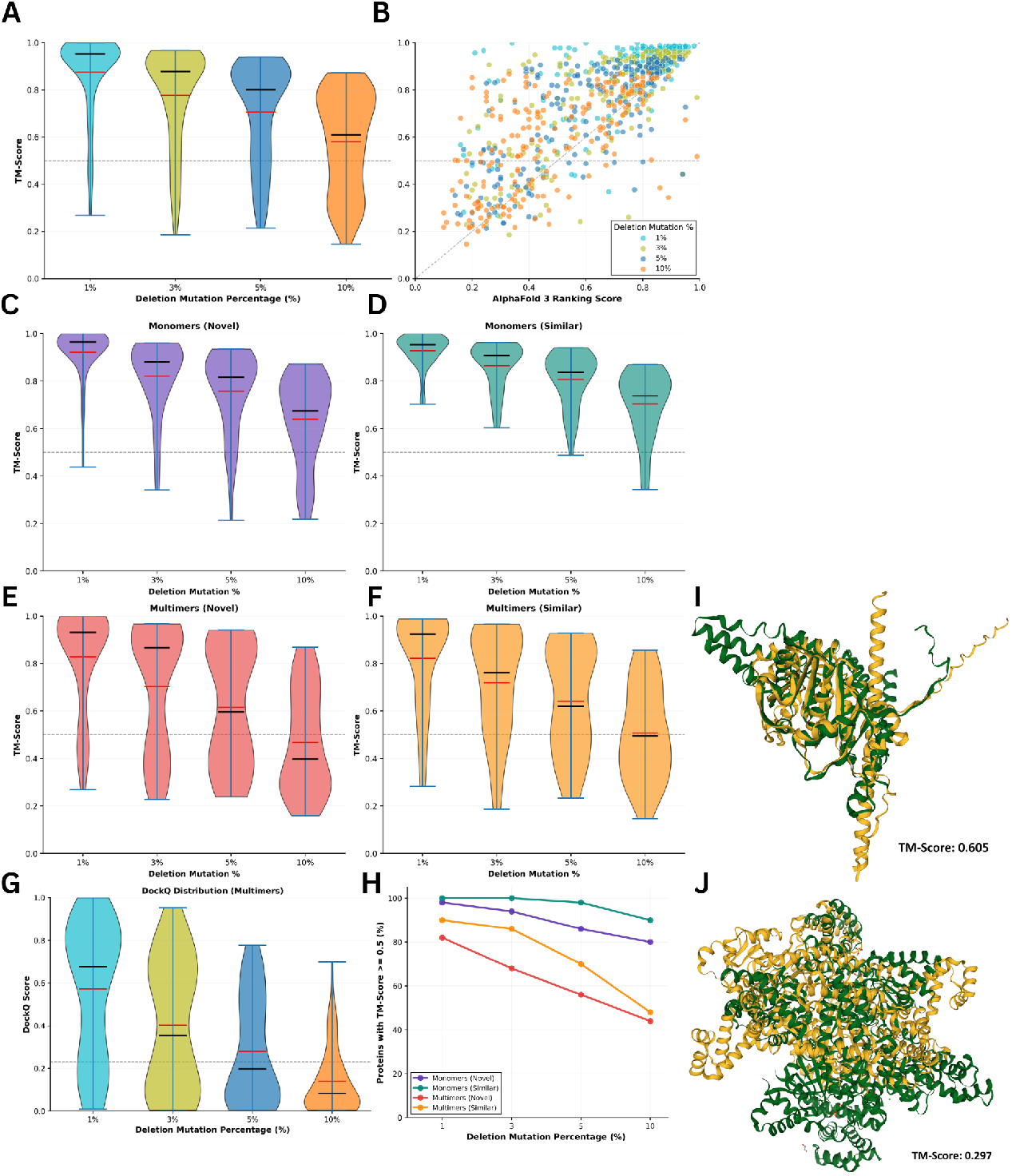
AlphaFold 3 exhibits structural invariance under adversarial deletion mutations. **A** Violin plots showing TM-score distributions for the full 200-protein dataset at each mutation threshold (1%, 3%, 5%, 10%), comparing mutated predictions to unmutated AlphaFold 3 predictions. **B** Correlation between AlphaFold 3 ranking confidence scores and structural accuracy (TM-score). Each point represents a single protein, colored by mutation percentage. **C–F** TM-score distributions stratified by protein category: (**C**) monomer-novel, (**D**) monomer-similar, (**E**) multimer-novel, and (**F**) multimer-similar. **G** DockQ score distributions for multimeric proteins (novel and similar bins combined), showing interfacial accuracy degradation across mutation thresholds. **H** Survival curve indicating the fraction of proteins maintaining accurate global fold (TM-score *≥* 0.5) at each mutation level. **I** Structural superposition of 8CQZ at 10% mutation (yellow) onto the unmutated prediction (green), showing preserved fold architecture (TM-score = 0.605). **J** Structural superposition of 7OCN at 10% mutation (yellow) onto the unmutated prediction (green), showing original fold destruction (TM-score = 0.297). All TM-scores and DockQ values reflect comparison to the unmutated AlphaFold 3 prediction rather than experimental structures.

The violin plots in Figure 2A and G illustrate TM-score and DockQ distributions across deletion thresholds. The relatively modest decrease in structural accuracy under deletion mutations—compared to point mutations—reflects the lower maximum mutation level tested. Nevertheless, the pattern of fold preservation is striking given that deletions near the protein core would be expected to cause substantial structural rearrangement or complete loss of fold in real proteins. The structural accuracy scores decline more steeply for multimeric proteins, crossing into low-confidence regimes by the 5% deletion threshold. Monomeric proteins, however, preserve high correspondence to the original protein structure across all tested deletion levels.

Furthermore, AlphaFold 3’s internal ranking score exhibit increased sensitivity under deletion-based perturbations, yet still display a notable degree of invariance relative to the severity of the structural disruption. The association between deletion fraction and ranking score is moderate, with a Pearson correlation coefficient of *−*0.514 and a Spearman rank correlation of *−* 0.491. The mean ranking score declines steadily as the proportion of deleted residues increases, decreasing from 0.67 at 1% deletions to 0.60 at 3%, 0.50 at 5%, and 0.32 at 10%. While this downward trend is more pronounced than in the point-mutation experiment, the model’s confidence remains relatively high at low deletion levels even thought sparse deletions—particularly within the protein core—would be expected to substantially impair structural integrity in real proteins [48, 49]. This persistence of elevated confidence scores mirrors the observed invariance in predicted structural accuracy.

### 2.3 Comparison of AlphaFold 3 and ESMFold Under Adversarial Mutation

To determine whether the structural invariance observed under mutation is specific to AlphaFold 3 or reflects a broader property of deep learning–based structure prediction, we compared AlphaFold 3 with ESMFold across the same mutational regimes. Because ESMFold is not designed to process multimeric inputs, this comparison was restricted to monomeric proteins, and AlphaFold 3’s monomeric predictions were used accordingly.

Figure 3 presents the results of this analysis. Figure 3A and Figure 3B show the individual trajectories of each protein in terms of TM-score accuracy under point mutations and deletion mutations, respectively. Under deletion mutations, ESMFold and AlphaFold 3 perform comparably, with ESMFold exhibiting consistently greater flexibility—that is, lower structural similarity to its original prediction—across all deletion thresholds. This suggests that ESMFold’s predictions are slightly more sensitive to the disruptive effects of residue removal.Whether they yield experimentally correct structures is still an open question.

**Fig. 3.**
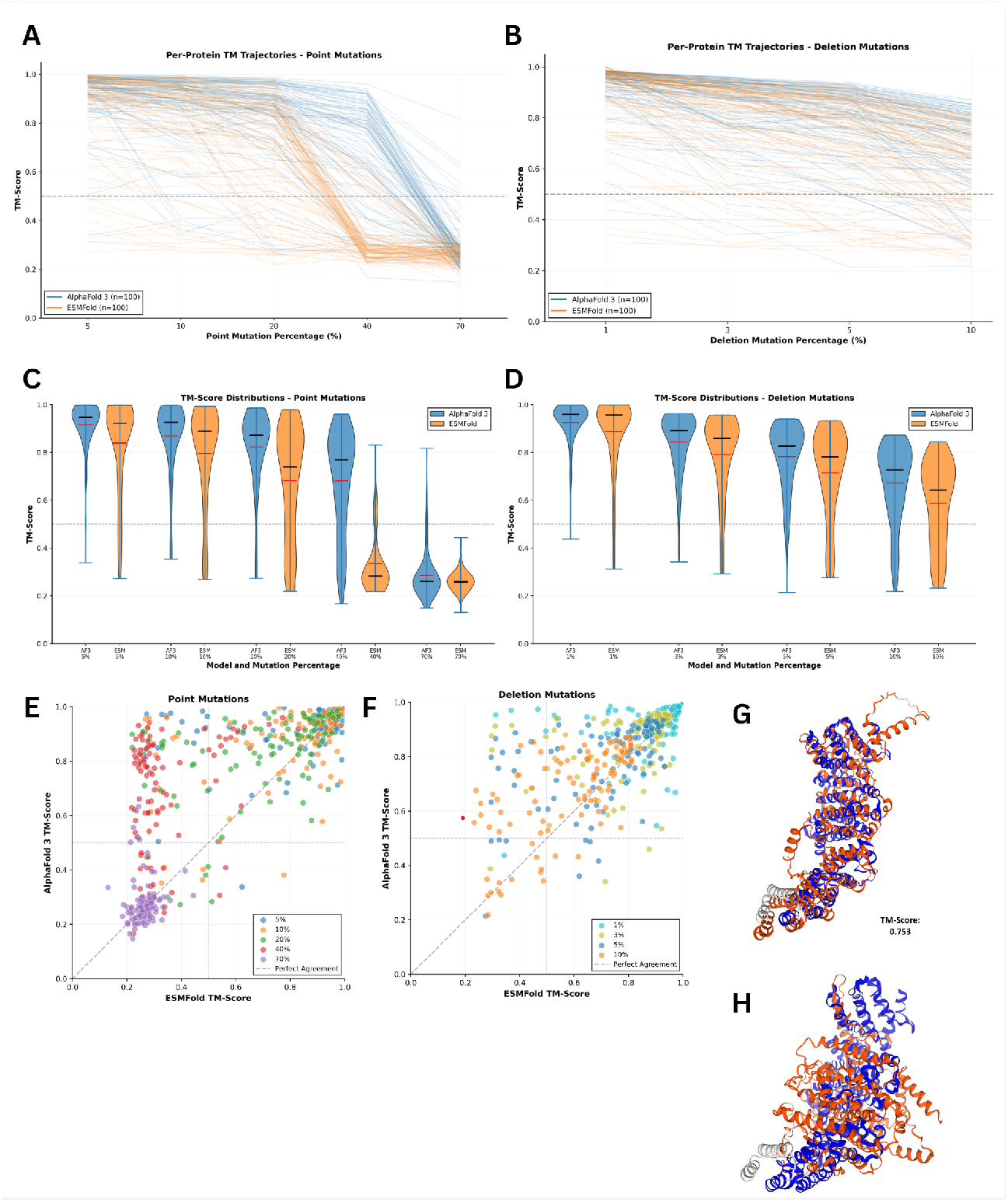
ESMFold exhibits greater sensitivity to point mutations than AlphaFold 3. **A** Individual protein accuracy trajectories under point mutations. Each line represents a single protein, with ESMFold (orange) and AlphaFold 3 (blue) predictions compared to their respective unmutated structures. **B** Individual protein accuracy trajectories under deletion mutations, colored as in **A. C** TM-score distributions across point mutation thresholds for ESMFold (orange) and AlphaFold 3 (blue). **D** TM-score distributions across deletion mutation thresholds, colored as in **C. E** Direct comparison of model accuracy under point mutations. Each point represents a protein at a given mutation threshold, with color indicating mutation percentage. Points above the diagonal indicate greater accuracy for AlphaFold 3. **F** Direct comparison of model accuracy under deletion mutations, displayed as in **E. G** Structural superposition of AlphaFold 3 predictions for protein 8RQT: 40% mutated structure (red) aligned to unmutated prediction (blue), demonstrating structural preservation (TM-score = 0.753). **H** Structural superposition of ESMFold predictions for protein 8RQT: 40% mutated structure (red) aligned to unmutated prediction (blue), demonstrating original fold destruction (TM-score = 0.293). ESMFold shows substantially greater structural divergence at intermediate mutation levels, suggesting higher sensitivity to sequence perturbations. All comparisons are between mutated and unmutated predictions from the same model.

The contrast is far more pronounced for point mutations, however. Between 20% and 40% point mutation, ESMFold’s accuracy with respect to its original predicted structure declines quickly and starkly, while AlphaFold 3 maintains its trend of gradual decrease. It is not until 70% mutation—corresponding to only 30% sequence identity—that AlphaFold 3 exhibits a comparable collapse, catching up to ESMFold’s degraded performance. This pattern is also evident in Figure 3E, a scatter plot demonstrating that ESMFold performance falls substantially below that of AlphaFold 3 at intermediate mutation levels.

We also applied ESMFold to the same fifteen fold-switching proteins analyzed with AlphaFold 3 (Supplementary Table S1). Consistent with the broader 200-protein dataset, ESMFold exhibited greater sensitivity to point mutations, with predicted structures diverging substantially from the original prediction beyond the 10% mutation threshold—though the effect was less pronounced than in the full dataset. These results reinforce the conclusion that ESMFold better captures the structural consequences of sequence perturbation, even for proteins known to undergo conformational switching.

These results indicate that ESMFold is more sensitive to deleterious point mutations and more attuned to their expected biophysical impact. While both models eventually converge at extreme mutation levels, ESMFold’s earlier response to sequence perturbation suggests it may better capture the relationship between sequence and structure, or, conversely, that AlphaFold 3’s invariance reflects a stronger reliance on learned templates that persist even when the underlying sequence has diverged substantially from biophysically viable configurations.

### 2.4 Evaluation of Confidence Metrics in AlphaFold 2 versus AlphaFold 3

To assess how reliably AlphaFold 2 and AlphaFold 3 estimate the accuracy of their own predictions, we evaluated both systems on one hundred proteins drawn from the monomer-novel and multimer-novel bins [1, 11]. This restriction avoids giving AlphaFold 3 an advantage from its broader training distribution and ensures a fair comparison [8]. Predicted structures were compared against experimentally determined structures from the Protein Data Bank, rather than against the original unmutated predictions used in the adversarial analyses above.

As shown in Figure 4, AlphaFold 3 achieves statistically significantly higher accuracy than AlphaFold 2, particularly for multimeric complexes. This improvement has been noted in prior work [8, 11], but another critical question is whether either model can reliably identify which of its generated predictions is most accurate. Standard practice is to generate five candidate structures and select the top-ranked prediction based on internal confidence metrics.

**Fig. 4.**
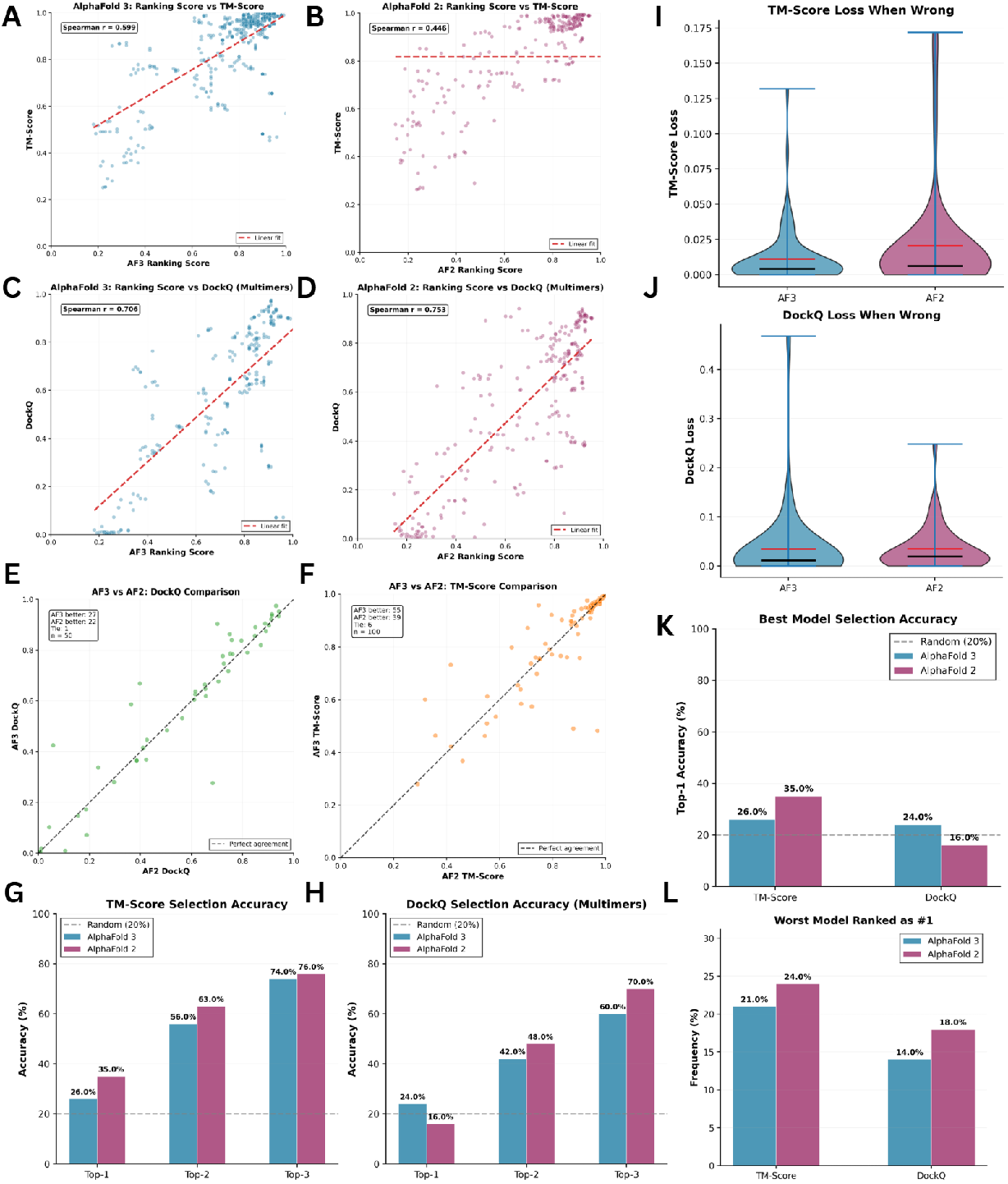
AlphaFold 2 and AlphaFold 3 confidence metrics fail to reliably identify the most accurate predictions. **A–D** Correlation between ranking confidence scores and structural accuracy for AlphaFold 3 (**A, C**) and AlphaFold 2 (**B, D**). TM-score is shown for monomers (**A, B**) and DockQ for multimers (**C, D**). Spearman correlation coefficients are indicated, with lines of best fit shown in red. **E–F** Direct comparison of structural accuracy between AlphaFold 2 and AlphaFold 3 on novel proteins, measured by (**E**) TM-score (monomers and multimers) and (**F**) DockQ (multimers only). Points above the diagonal indicate superior AlphaFold 3 performance. **G** Frequency with which each model selects the most accurate structure (rank 1), or a structure within the top 2 or top 3 most accurate, based on TM-score across 100 proteins. AlphaFold 3 (blue) and AlphaFold 2 (purple). **H** Model selection accuracy based on DockQ for 50 multimeric proteins, displayed as in **G. I** Distribution of TM-score loss incurred when the model-selected structure is not the most accurate prediction. Larger values indicate more severe selection errors. **J** Distribution of DockQ loss incurred by incorrect model selection, displayed as in **I. K** Combined view of model selection accuracy for both TM-score and DockQ metrics, showing the percentage of proteins for which each model correctly identifies the best prediction. **L** Frequency with which each model selects the least accurate (worst) prediction, highlighting severe ranking failures. Both models exhibit poor calibration, selecting the optimal structure in only 16–35% of cases.

Figure 4K reveals the limitations of both models’ confidence-based selection. AlphaFold 3 selects the most accurate structure—by TM-score or DockQ—only approximately 25% of the time, while AlphaFold 2 achieves at most 35%. Even when the definition of success is relaxed to include selecting among the top two or three most accurate structures, as shown in Figure 4G and Figure 4H, accuracy increases but never exceeds 80% for either model.

Incorrect model selection is not inconsequential. On average, selecting the wrong model costs approximately 0.01 TM-score points for AlphaFold 3 and 0.02 for AlphaFold 2. DockQ loss averages approximately 0.03 for both models. However, in some instances the losses are far more severe: up to 0.17 for TM-score and over 0.40 for DockQ. Figure 4I and Figure 4J illustrate that choosing the wrong model is not always a harmless error—–substantial accuracy can be forfeited when confidence metrics fail to identify the best prediction.

We also evaluated the correlation between each model’s ranking score—a linear combination of ipTM if applicable, pTM and clash penalties—and structural accuracy against the ground truth. The Spearman correlation coefficient was higher for AlphaFold 2 on multimeric proteins but higher for AlphaFold 3 on monomeric proteins, as shown in Figure 4A–D. The linear relationship between predicted score and accuracy, quantified by the Pearson correlation coefficient, was stronger for multimers than for monomers in both models. For multimers, the correlations were 0.72 for AlphaFold 3 and 0.83 for AlphaFold 2, indicating greater sensitivity of AlphaFold 2’s ranking score in this setting. In contrast, for monomers the correlations were 0.59 for AlphaFold 3 and 0.44 for AlphaFold 2, suggesting superior performance of AlphaFold on monomeric predictions. These results indicate that while confidence metrics provide useful signal, they remain imperfect guides for selecting the most accurate prediction—a limitation with practical implications for downstream applications that depend on model-selected structures.

### 2.5 Structural Template Availability Predicts AlphaFold 3 Confidence

To shed light on what AlphaFold 3’s confidence metrics truly measure, we explored whether they are correlated with the availability of structurally similar training-set templates, we performed an exhaustive structural search of the pre-cutoff Protein Data Bank using FoldSeek [50]. For those proteins in the 200-constituent protein dataset with relevant templates, we identified the most structurally similar chain deposited on or before the AlphaFold 3 training cutoff and quantified its relationship to model confidence (See Methods for more details.)

AlphaFold 3’s ranking score correlates significantly with the query TM-score of the best pre-cutoff structural template (Pearson *r* = 0.39, *p* = 7.8 *×* 10^*−*5^; Spearman *ρ* = 0.39, *p* = 8.2 *×* 10^*−*5^). By contrast, sequence identity to that same template is a weak and non-significant predictor (*r* = 0.033, *p* = 0.744), indicating that confidence tracks structural rather than sequence similarity to training-set exemplars. A comparable, though weaker, effect holds for multimers (*r* = 0.27, *p* = 7.7 *×* 10^*−*3^; *ρ* = 0.30, *p* = 3.1 *×* 10^*−*3^).

Stratifying monomers by novelty reveals that the correlation is robust regardless of whether the query sequence appears in the training distribution—novel proteins (*r* = 0.51, *p* = 1.9 *×* 10^*−*4^) and in-dataset proteins (*r* = 0.49, *p* = 3.1 *×* 10^*−*4^) show comparable effect sizes despite differing substantially in their template landscapes (Supplementary Table S4). For multimers, the correlation is significant among novel complexes (*r* = 0.43, *p* = 2.2 *×* 10^*−*3^) but absent for in-dataset complexes (*r* = 0.14, *p* = 0.34), suggesting sequence memorization supplants template dependence when the query falls within the training distribution.

For monomers, the single best structural template is the strongest predictor of confidence (Spearman *ρ* = 0.35, *p* = 4.4 *×* 10^*−*4^), with correlations declining for deeper neighborhoods and neighborhood density offering no additional predictive value. Multimers exhibit the opposite trend: correlations increase monotonically from *ρ* = 0.30 at the top hit to *ρ* = 0.46 at the top 100 hits (*p* = 2.3 *×* 10^*−*6^), suggesting that for complex prediction, breadth of structural coverage in the training set matters more than any single exemplar. See Supplementary Table S4 for more details.

## 3 Discussion

The point and deletion mutation analyses presented here reveal a striking invariance in both the structures predicted by AlphaFold 3 and in its confidence metrics under substantial sequence perturbations. Using the commonly accepted TM-score threshold of 0.5 to indicate similar folds [43], we observed that up to 40 percent of residues in a protein can be mutated—and up to 10 percent of residues deleted—before the predicted structure meaningfully diverges from the original AlphaFold 3 prediction on average. This trend persists even for fold-switching proteins, which are experimentally known to adopt multiple conformations, highlighting a verifiable disconnect between model predictions and expected biological behavior [45].

These observations have important implications for how the field interprets and applies structure prediction models. The degree of invariance we document suggests that AlphaFold 3 may not be processing input sequences at the granularity required to capture the structural consequences of individual residue changes. While previous work has noted that AlphaFold exhibits some insensitivity to point mutations, the extent of this phenomenon warrants far closer examination [30]. Insensitivity to one or two amino acid substitutions might be expected, as such minor perturbations may genuinely leave the global fold intact in many cases. However, changing up to 70% of the amino acids in an intentionally deleterious manner, to the point where the protein shares minimal sequence identity with the original, while still observing the same predicted global fold raises questions about the nature of the learned sequence-structure mapping.

One interpretation of these results is that AlphaFold 3 relies heavily on identifying similarities to known templates in its training data rather than reasoning about sequence-structure relationships from biophysical principles, which is consistent with studies showing that AlphaFold over-relies on its training set when predicting alternative conformations [35]. Thus caution should be exercised when it is a major component in any protein design algorithm.

The comparison with ESMFold provides additional context. We find that ESMFold exhibits greater sensitivity to point mutations than AlphaFold 3, with predicted structures diverging more rapidly as mutation load increases. This observation aligns with research demonstrating that ESM models, trained with masked language modeling objectives, learn coevolutionary statistics and comparative relationships between amino acids [51]. The self-supervised training paradigm forces these models to somewhat internalize which residues are plausible in a given sequence context, implicitly encoding information about substitution tolerability. While ESMFold achieves lower absolute accuracy for structure prediction than AlphaFold 3, its greater responsiveness to sequence perturbation may reflect a different—and in some contexts more appropriate—learned representation of sequence–structure relationships, a property that is also evident when predicting fold-switching proteins, where ESMFold displays increased sensitivity to mutations known to induce conformational transitions.

These findings also underscore concerns about confidence metric reliability. Our analysis shows that neither AlphaFold 2 nor AlphaFold 3 reliably identifies the most accurate structure among five generated predictions, with both models selecting the best structure only approximately 16–35% of the time. Previous studies have documented issues with AlphaFold confidence metrics [38, 52–54], and recent work has shown that modifications to confidence metric calculations do not fully address the underlying bias in the data produced by the model [8]. AlphaFold 3 employs a more sophisticated atomic-level confidence module and benefits from a broader training distribution than AlphaFold 2 [1, 11], yet the differences in model selection accuracy between versions are modest. This suggests that the limitation may be fundamental to how these models learn to estimate their own uncertainty rather than a matter of implementation detail. The structural template analysis provides quantitative support for this: AlphaFold 3’s confidence correlates significantly with the structural quality of the best pre-cutoff template identified by FoldSeek, suggesting that the model’s uncertainty estimates reflect template availability more than genuine assessment of prediction quality.

The practical consequences merit consideration. When an incorrect model is selected, the losses in accuracy can be substantial—reaching up to 0.17 in TM-score and over 0.4 in DockQ in some cases. For applications in which AlphaFold predictions inform downstream decisions—whether in drug discovery, protein engineering, or pathway analysis—these errors could lead to the exclusion of promising candidates or the retention of problematic ones [14, 17]. If these models are insensitive to certain classes of mutations, design pipelines built upon them may inherit analogous limitations, potentially generating implausible sequences or failing to explore viable regions of sequence space.

These observations, however, do not diminish the transformative impact of AlphaFold on structural biology. For the majority of proteins and applications, AlphaFold 3 achieves remarkable accuracy and has enabled research that would otherwise be impractical. The limitations we document are most relevant for out-of-distribution cases and for applications requiring fine-grained sensitivity to sequence variation.

The source of these limitations remains an open question. One possibility is architectural: the transformer and diffusion-based frameworks underlying current structure prediction models may inherently favor interpolation within the training distribution over extrapolation beyond it. Another possibility lies in the training data itself. The structural databases on which these models are trained consist almost exclusively of stable, wild-type proteins—mutant proteins, particularly those carrying destabilizing mutations that alter or abolish the native fold, are largely absent. Without exposure to examples of mutation-induced structural disruption, models may never learn the relationships necessary to predict such effects. ESMFold’s greater sensitivity to mutations is consistent with this view: its masked language modeling objective, which requires predicting plausible residues from sequence context, implicitly encodes information about which substitutions are and are not tolerable. A third possibility is that these models operate within an inherent radius of convergence, and that only at higher mutation counts does the structural perturbation become large enough for the model to respond meaningfully.

The ecosystem of tools built upon AlphaFold amplifies the importance of understanding its limitations. Many state-of-the-art methods for structural prediction, protein design and protein-protein interaction analysis incorporate AlphaFold’s architecture or use its predictions as inputs [14, 17]. If the underlying model harbors systematic biases, these may propagate through derivative tools. This is not merely a theoretical concern—it has practical implications for drug discovery, protein design and its application to the understanding of protein evolution.

Several directions may help address the limitations we describe. Training on datasets that include destabilized mutants and misfolded proteins could expose models to a broader range of sequence-structure relationships. Incorporating explicit biophysical priors—such as energy functions or molecular dynamics data—into model architectures or loss functions may help enforce realistic structural responses to mutation. Developing improved confidence estimation methods that better calibrate uncertainty, particularly for out-of-distribution inputs, would enhance the reliability of model-guided decisions. Finally, continued benchmarking against adversarial and edge cases will be essential for characterizing the boundaries of model applicability.

The remarkable successes of AlphaFold should not obscure the work that remains. Structure prediction has advanced enormously, but the ability to reliably predict how arbitrary proteins—including those with extensive mutations or novel sequences— will fold remains incomplete. Recognizing the current limitations of these tools is a necessary step toward developing the next generation of methods that more fully capture the relationship between protein sequence and structure.

## 4 Methods

### 4.1 AlphaFold Testing Dataset Collection

To rigorously evaluate AlphaFold’s performance, we assembled a dataset of 200 proteins, divided evenly into four categories: multimer-novel, multimer-similar, monomer-novel, and monomer-similar, with 50 proteins per category. “Multimeric” and “monomeric” indicate whether a protein contains multiple chains or a single chain, respectively, while “novel” and “similar” denote sequence similarity to proteins in the PDB released on or before September 30, 2021 [11, 55]. Proteins labeled as similar share 30% or more sequence homology with pre-September 2021 PDB entries, meaning their homologs could plausibly have been in AlphaFold 3’s training set [8]. In contrast, novel proteins have no such homology. Importantly, all proteins in the dataset were released after September 30, 2021, and no two proteins share 30% or more sequence similarity, ensuring minimal redundancy and that none were present in AlphaFold 3’s training set. This categorization enables a detailed assessment of AlphaFold’s performance across different protein types and levels of novelty.

### 4.2 Fold-Switching Protein Dataset

To extend the mutational analysis to proteins with experimentally validated alternative conformations, we selected fifteen fold-switching proteins from the curated dataset compiled by Porter et al. [45]. All proteins in this set are monomeric and have been experimentally confirmed to adopt multiple distinct folds under different conditions. The selected proteins correspond to the following PDB entries: 2KXO, 2LSH, 4OV8, 2MZ7, 4PMK, 2N4O, 2KTM, 2LE3, 2X9C, 3J9E, 5SUZ, 4HLS, 1S5P, 3TKA, and 3GAX. For these proteins, only chain A from their corresponding PDB files was used, as described in [45]. Point mutations were introduced to these proteins using procedures similar to those applied to the 200-protein dataset described above, with the exception of a modified position-weighting scheme that prioritized residues or regions known to induce fold-switching. Details of this scheme are provided in the Supplementary Table S3. MSA processing and AlphaFold 3 inference followed the same protocols, with templates excluded. This analysis allowed us to assess whether proteins known to exhibit conformational plasticity show greater structural responsiveness to mutations in AlphaFold 3’s predictions compared to proteins without documented fold-switching behavior.

### 4.3 Mutation Strategy

The mutation percentages applied differed between point mutations and deletion mutations to reflect their relative biophysical severity. Point mutations were tested at 5%, 10%, 20%, 40%, and 70% thresholds, while deletion mutations were tested at 1%, 3%, 5%, and 10% thresholds. This asymmetry reflects the fact that deletion mutations are substantially more deleterious than point mutations [46]. Even a single residue deletion in a critical region can cause severe structural destabilization or complete loss of fold, whereas point mutations—particularly conservative substitutions—are often tolerated without catastrophic consequences [47]. By using lower deletion percentages, we ensured that the perturbations remained within a regime where structural prediction could be meaningfully evaluated, while still probing the model’s sensitivity to this severe class of mutation.

Mutations were applied cumulatively across all thresholds. Each higher mutation percentage retained all mutations from the previous level and added new mutations to reach the target threshold. For example, the 20% point mutation configuration contained all mutations present in the 10% configuration, plus additional mutations to reach 20% of the sequence length. Similarly, the 5% deletion configuration included all deletions from the 3% level, supplemented with further deletions. This cumulative strategy ensures that differences in structural predictions across mutation levels reflect the incremental addition of perturbations rather than independent sampling at each threshold, enabling a more controlled assessment of mutational sensitivity.

For homomeric complexes, mutations were applied synchronously: the same residue positions were mutated to the same amino acids across all identical chains. For example, in a homodimeric protein, if position 42 was mutated from serine to tryptophan in chain A, position 42 was also mutated from serine to tryptophan in chain B. This approach reflects the biological reality that homomeric complexes consist of identical gene products and thus share identical sequences. In contrast, heteromeric complexes have distinct chains with independent sequences, and mutations were applied independently to each chain, such that different positions and amino acid substitutions were selected for each unique chain type.

### 4.4 AlphaFold Model Configurations and Inputs

For all mutational analyses, AlphaFold 3 was run using its default configuration, with ten recycling steps and five diffusion samples [11]. The primary modifications were applied to the input features. Template models—which AlphaFold 3 uses to infer general structural information and which could artificially inflate mutational invariance—were excluded [11]. For both deletion and point mutations, the same random seed was used across all mutation thresholds to ensure comparability of model accuracy. All default settings were used for MSA generation in AlphaFold 3.

In the deletion mutation regime, multiple sequence alignments (MSAs) were modified by removing columns corresponding to deleted residues (see Algorithm 2 for pseudocode). This ensured compatibility with AlphaFold 3, which cannot process MSAs containing sequences whose lengths differ from that of the input sequence. In contrast, for the point mutation regime, MSAs were left unchanged and only the input sequences were mutated, leveraging the fact that MSAs naturally encode organismal sequence variation (see Algorithm 1 for pseudocode).

For deletion mutations, the paired MSA was excluded from the input data. Previous studies have shown that paired MSAs do not improve structural prediction accuracy; thus, they were omitted to simplify the required modifications.

#### Algorithm 1

MSA Processing for Point Mutations

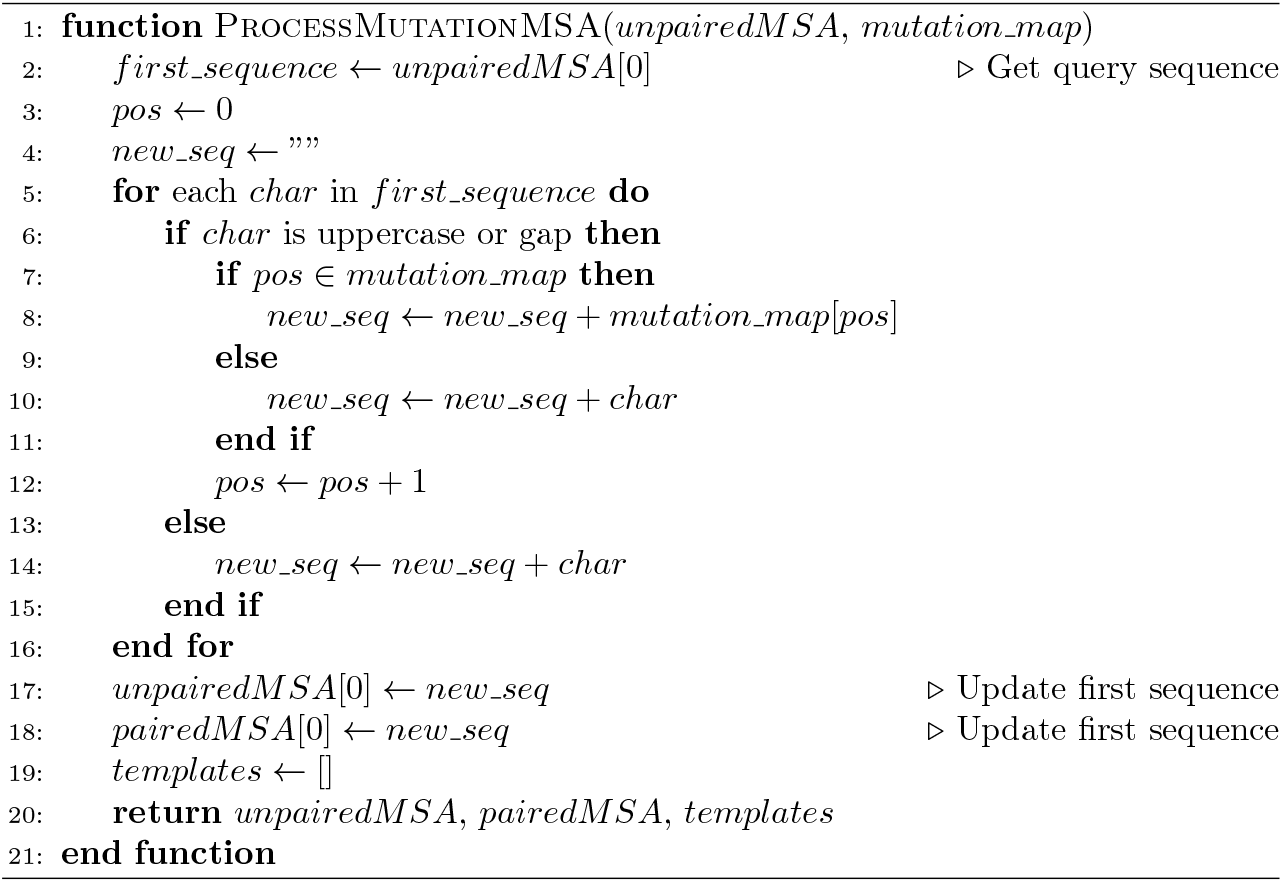

#### Algorithm 2

MSA Processing for Deletion Mutations

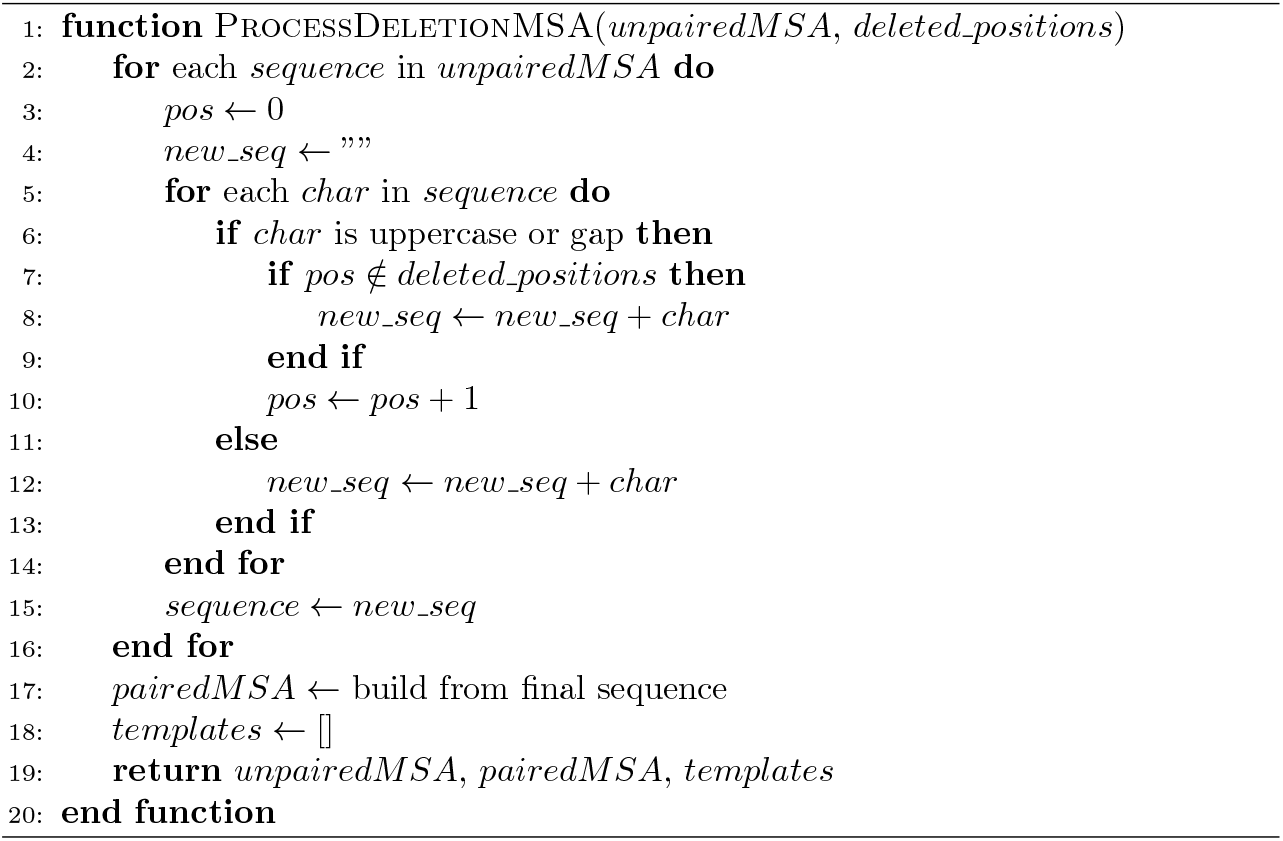

To verify that modifying MSAs—rather than regenerating them after mutation— did not introduce artificial mutational invariance, MSAs were fully regenerated for the same 200 proteins in the dataset using AlphaFold 3’s standard MSA generation pipeline. These proteins were first mutated at all tested thresholds (5%, 10%, 20%, 40%, and 70% for point mutations; 1%, 3%, 5%, and 10% for deletions), and then new MSAs were generated de novo for each mutated sequence using the default AlphaFold 3 MSA generation pipeline. The paired MSA was retained in this regeneration experiment, as it was accurately generated for the mutated sequences. No structural templates were used. No major differences in TM-score or AlphaFold 3 ranking score between models using modified versus regenerated MSAs were noted. Interestingly, models with regenerated MSAs exhibited slightly greater mutational invariance than their directly modified counterparts, suggesting that MSA regeneration only exacerbates the model’s structural invariance.

For the comparative analysis between AlphaFold 3 and AlphaFold 2, both models were configured with ten recycles and five samples [1, 11]. AlphaFold 3 was run without templates to ensure that its template-based features, which are absent in AlphaFold 2, did not confer an unfair performance advantage. As noted in the Results section, this analysis was conducted on 100 proteins drawn from novel bins—50 monomers and 50 multimers—which had no homologs in the AlphaFold 3 training set.

### 4.5 Protein Mutation Details

The point mutations applied to proteins in the 200-protein dataset were designed to be as deleterious as possible, ensuring that the mutations were unlikely to incidentally enhance structural stability. The mapping of these mutations is summarized in the Table S2 in the Supplementary Information. To further increase the likelihood of major structural disruption, the residue mutation algorithms for both point and deletion mutations incorporated a positional bias that preferentially selected residues near the center of the protein [40]. This bias was implemented mathematically by assigning each residue *i* a weight

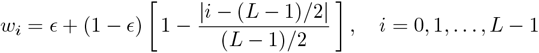

where *L* is the sequence length and *ϵ* defines the weight for residues at the sequence termini, which, for the purposes of the study, was set to 0.1. This scheme ensures that central residues, which are more likely to contribute to the protein’s core stability, have a higher probability of mutation.

### 4.6 ESMFold Configuration and Analysis

To compare AlphaFold 3’s mutational sensitivity with that of an alternative deep learning–based structure prediction model, we performed parallel analyses using ESMFold [39, 51]. The ESMFold implementation was obtained from the *HuggingFace* [56, 57] repository and run with all default configuration settings. Because ESMFold was not trained to predict multimeric protein structures, this comparison was restricted to monomeric proteins only [39]. Specifically, we analyzed the 100 monomeric proteins from the 200-protein dataset, comprising 50 from the monomer-novel bin and 50 from the monomer-similar bin. Point mutations and deletion mutations were applied to these proteins using the identical mutation strategy employed for AlphaFold 3. For each mutated protein, ESMFold predictions were compared to the original unmutated ESMFold prediction.

### 4.7 FoldSeek Exhaustive Structural Search

Each of the 200 experimentally determined structures was queried against FoldSeek’s PDB100 database [50], which comprises approximately 340,000 representative chains clustered at 100% sequence identity from the full Protein Data Bank. Searches were conducted in exhaustive mode (`--exhaustive-search 1`) with TM-align-based scoring (`--alignment-type 1`), bypassing FoldSeek’s heuristic prefilter to guarantee globally optimal structural matches, yielding approximately 9,800 hits per query. Results were post-filtered to retain only hits deposited on or before September 30, 2021, using a list of approximately 180,000 pre-cutoff PDB identifiers obtained from RCSB PDB [55]. The query-normalized TM-score (qtmscore) was used as the primary structural similarity metric, as normalizing by query length prevents partial domain matches from inflating similarity estimates.

For monomeric proteins, AlphaFold 3’s monomeric ranking score, pTM, was used as the confidence metric; for multimeric complexes, the ranking score (0.8 *×* ipTM + 0.2 *×* pTM) was used [11], jointly capturing fold accuracy and interface quality. For monomers, the top 100 pre-cutoff hits were extracted per query and ranked by qtm-score. One monomer had no pre-cutoff hits, yielding *n* = 99 for the primary correlation analyses; three further monomers had fewer than 100 pre-cutoff hits, yielding *n* = 96 for the neighborhood density analysis. For multimeric complexes, FoldSeek’s monomer search mode decomposed each complex into constituent chains, which were searched independently. Per-chain neighborhood statistics were computed from the top 100 pre-cutoff hits and averaged across chains to obtain complex-level estimates. Two multimers were excluded because at least one chain had fewer than 100 pre-cutoff hits, yielding *n* = 98 used throughout. Minimum-across-chains and chain-length-weighted aggregation schemes were also tested and produced comparable results. Only mean aggregation is reported for simplicity.

### 4.8 Structure Comparison Metrics

TM-score and DockQ were calculated using the OpenStructure package [58]. TM-score quantifies global structural similarity between two protein structures and is normalized to a range of 0 to 1, where values above 0.5 typically indicate the same fold [42]. DockQ assesses the quality of predicted protein-protein interfaces by combining information about interface RMSD, ligand RMSD, and native contact fraction [44]. For multimeric proteins with more than two chains, the weighted average DockQ score across all pairwise chain interfaces was computed and used as the reported metric, with a standard threshold of 0.23 used for a valid interfacial conformation [44, 59]. Both metrics were calculated by comparing mutated predictions to the original unmutated AlphaFold 3 prediction for the mutational invariance analyses, or to experimental structures from the Protein Data Bank for the model comparison analyses.

## Acknowledgments

This research was supported in part by grant GM-118039 from the Division of General Medical Sciences of the National Institutes of Health . We thank Jessica Forness for proofreading and polishing this manuscript.

## References

[1] Jumper, J., Evans, R., Pritzel, A., Green, T., Figurnov, M., Ronneberger, O., Tunyasuvunakool, K., Bates, R., Žídek, A., Potapenko, A., Bridgland, A., Meyer, C., Kohl, S.A.A., Ballard, A.J., Cowie, A., Romera-Paredes, B., Nikolov, S., Jain, R., Adler, J., Back, T., Petersen, S., Reiman, D., Clancy, E., Zielinski, M., Steinegger, M., Pacholska, M., Berghammer, T., Bodenstein, S., Silver, D., Vinyals, O., Senior, A.W., Kavukcuoglu, K., Kohli, P., Hassabis, D.: Highly accurate protein structure prediction with alphafold. Nature 596(7873), 583–589 (2021) 10.1038/s41586-021-03819-2. Accessed 2024-08-31

[2] Bertoline, L.M.F., Lima, A.N., Krieger, J.E., Teixeira, S.K.: Before and after alphafold2: An overview of protein structure prediction. Frontiers in Bioinformatics 3, 1120370 (2023) 10.3389/fbinf.2023.1120370. Accessed 2025-03-12

[3] Bryant, P., Pozzati, G., Elofsson, A.: Improved prediction of protein-protein interactions using alphafold2. Nature Communications 13(1), 1265 (2022) 10.1038/s41467-022-28865-w. Publisher: Nature Publishing Group. Accessed 2024-09-14

[4] Evans, R., O’Neill, M., Pritzel, A., Antropova, N., Senior, A., Green, T., Žídek, A., Bates, R., Blackwell, S., Yim, J., Ronneberger, O., Bodenstein, S., Zielinski, M., Bridgland, A., Potapenko, A., Cowie, A., Tunyasuvunakool, K., Jain, R., Clancy, E., Kohli, P., Jumper, J., Hassabis, D.: Protein complex prediction with AlphaFold-Multimer. bioRxiv. Pages: 2021.10.04.463034 Section: New Results (2022). 10.1101/2021.10.04.463034. https://www.biorxiv.org/content/10.1101/2021.10.04.463034v2 Accessed 2024-09-15

[5] Tunyasuvunakool, K., Adler, J., Wu, Z., Green, T., Zielinski, M., Žídek, A., Bridgland, A., Cowie, A., Meyer, C., Laydon, A., Velankar, S., Kleywegt, G.J., Bateman, A., Evans, R., Pritzel, A., Figurnov, M., Ronneberger, O., Bates, R., Kohl, S.A.A., Potapenko, A., Ballard, A.J., Romera-Paredes, B., Nikolov, S., Jain, R., Clancy, E., Reiman, D., Petersen, S., Senior, A.W., Kavukcuoglu, K., Birney, E., Kohli, P., Jumper, J., Hassabis, D.: Highly accurate protein structure prediction for the human proteome. Nature 596(7873), 590–596 (2021) 10.1038/s41586-021-03828-1

[6] Tsitsa, I., Conev, A., David, A., Islam, S.A., Sternberg, M.J.E.: The alphafold database ages. bioRxiv (2025) 10.1101/2025.06.22.660930 https://www.biorxiv.org/content/early/2025/09/27/2025.06.22.660930.full.pdf

[7] Kovalevskiy, O., Mateos-Garcia, J., Tunyasuvunakool, K.: Alphafold two years on: Validation and impact. Proceedings of the National Academy of Sciences 121(34), 2315002121 (2024) 10.1073/pnas.2315002121 https://www.pnas.org/doi/pdf/10.1073/pnas.2315002121

[8] Feldman, J., Skolnick, J.: AF3Complex Yields Improved Structural Predictions of Protein Complexes. bioRxiv. Pages: 2025.02.27.640585 Section: New Results (2025). 10.1101/2025.02.27.640585. https://www.biorxiv.org/content/10.1101/2025.02.27.640585v1 Accessed 2025-03-12

[9] Gao, M., Nakajima An, D., Parks, J.M., Skolnick, J.: Af2complex predicts direct physical interactions in multimeric proteins with deep learning. Nature Communications 13(1), 1744 (2022) 10.1038/s41467-022-29394-2. Accessed 2024-09-14

[10] Gao, M., Skolnick, J.: Improved deep learning prediction of antigen-antibody interactions. Proceedings of the National Academy of Sciences of the United States of America 121(41), 2410529121 (2024) 10.1073/pnas.2410529121. Accessed 2025-02-17

[11] Abramson, J., Adler, J., Dunger, J., Evans, R., Green, T., Pritzel, A., Ronneberger, O., Willmore, L., Ballard, A.J., Bambrick, J., Bodenstein, S.W., Evans, D.A., Hung, C.-C., O’Neill, M., Reiman, D., Tunyasuvunakool, K., Wu, Z., Žemgulytė, A.e., Arvaniti, E., Beattie, C., Bertolli, O., Bridgland, A., Cherepanov, A., Congreve, M., Cowen-Rivers, A.I., Cowie, A., Figurnov, M., Fuchs, F.B., Gladman, H., Jain, R., Khan, Y.A., Low, C.M.R., Perlin, K., Potapenko, A., Savy, P., Singh, S., Stecula, A., Thillaisundaram, A., Tong, C., Yakneen, S., Zhong, E.D., Zielinski, M., Žídek, A., Bapst, V., Kohli, P., Jaderberg, M., Hassabis, D., Jumper, J.M.: Accurate structure prediction of biomolecular interactions with alphafold 3. Nature 630(8016), 493–500 (2024) 10.1038/s41586-024-07487-w. Publisher: Nature Publishing Group. Accessed 2024-09-08

[12] Roy, R., Al-Hashimi, H.M.: Alphafold3 takes a step toward decoding molecular behavior and biological computation. Nature Structural & Molecular Biology 31(7), 997–1000 (2024) 10.1038/s41594-024-01350-2. Publisher: Nature Publishing Group. Accessed 2024-09-01

[13] Thompson, B., Petrić Howe, N.: Alphafold 3.0: the ai protein predictor gets an upgrade. Nature (2024) 10.1038/d41586-024-01385-x. Bandiera abtest: a Cg type: Nature Podcast Publisher: Nature Publishing Group Subject term: Exoplanets, Physics, Proteomics, Materials science. Accessed 2024-09-01

[14] Watson, J.L., Juergens, D., Bennett, N.R., Trippe, B.L., Yim, J., Eisenach, H.E., Ahern, W., Borst, A.J., Ragotte, R.J., Milles, L.F., Wicky, B.I.M., Hanikel, N., Pellock, S.J., Courbet, A., Sheffler, W., Wang, J., Venkatesh, P., Sappington, I., Torres, S.V., Lauko, A., De Bortoli, V., Mathieu, E., Ovchinnikov, S., Barzilay, R., Jaakkola, T.S., DiMaio, F., Baek, M., Baker, D.: De novo design of protein structure and function with rfdiffusion. Nature 620(7976), 1089–1100 (2023) 10.1038/s41586-023-06415-8. Publisher: Nature Publishing Group. Accessed 2025-07-21

[15] Yang, Z., Zeng, X., Zhao, Y., Chen, R.: Alphafold2 and its applications in the fields of biology and medicine. Signal Transduction and Targeted Therapy 8(1), 1–14 (2023) 10.1038/s41392-023-01381-z. Publisher: Nature Publishing Group. Accessed 2024-08-31

[16] Dauparas, J., Anishchenko, I., Bennett, N., Bai, H., Ragotte, R.J., Milles, L.F., Wicky, B.I.M., Courbet, A., Haas, R.J., Bethel, N., Leung, P.J.Y., Huddy, T.F., Pellock, S., Tischer, D., Chan, F., Koepnick, B., Nguyen, H., Kang, A., Sankaran, B., Bera, A.K., King, N.P., Baker, D.: Robust deep learning based protein sequence design using proteinmpnn. bioRxiv (2022) 10.1101/2022.06.03.494563 https://www.biorxiv.org/content/early/2022/06/04/2022.06.03.494563.full.pdf

[17] Stark, H., Faltings, F., Choi, M., Xie, Y., Hur, E., O’Donnell, T., Bushuiev, A., Uçar, T., Passaro, S., Mao, W., Reveiz, M., Bushuiev, R., Pluskal, T., Sivic, J., Kreis, K., Vahdat, A., Ray, S., Goldstein, J.T., Savinov, A., Hambalek, J.A., Gupta, A., Taquiri-Diaz, D.A., Zhang, Y., Hatstat, A.K., Arada, A., Kim, N.H., Tackie-Yarboi, E., Boselli, D., Schnaider, L., Liu, C.C., Li, G.-W., Hnisz, D., Sabatini, D.M., DeGrado, W.F., Wohlwend, J., Corso, G., Barzilay, R., Jaakkola, T.: Boltzgen: Toward universal binder design. bioRxiv (2025) 10.1101/2025.11.20.689494 https://www.biorxiv.org/content/early/2025/11/24/2025.11.20.689494.full.pdf

[18] Wohlwend, J., Corso, G., Passaro, S., Reveiz, M., Leidal, K., Swiderski, W., Portnoi, T., Chinn, I., Silterra, J., Jaakkola, T., Barzilay, R.: Boltz-1 Democratizing Biomolecular Interaction Modeling. bioRxiv. Pages: 2024.11.19.624167 Section: New Results (2024). 10.1101/2024.11.19.624167. https://www.biorxiv.org/content/10.1101/2024.11.19.624167v1 Accessed 2025-10-07

[19] Pacesa, M., Nickel, L., Schmidt, J., Pyatova, E., Schellhaas, C., Kissling, L., Alcaraz-Serna, A., Cho, Y., Ghamary, K.H., Vinué, L., Yachnin, B.J., Wollacott, A.M., Buckley, S., Georgeon, S., Goverde, C.A., Hatzopoulos, G.N., Gönczy, P., Muller, Y.D., Schwank, G., Ovchinnikov, S., Correia, B.E.: Bindcraft: one-shot design of functional protein binders. bioRxiv (2024) 10.1101/2024.09.30.615802 https://www.biorxiv.org/content/early/2024/10/01/2024.09.30.615802.full.pdf

[20] Zhang, Z., Wayment-Steele, H.K., Brixi, G., Wang, H., Kern, D., Ovchinnikov, S.: Protein language models learn evolutionary statistics of interacting sequence motifs. Proceedings of the National Academy of Sciences 121(45), 2406285121 (2024) 10.1073/pnas.2406285121. Publisher: Proceedings of the National Academy of Sciences. Accessed 2025-05-09

[21] Brixi, G., Durrant, M.G., Ku, J., Poli, M., Brockman, G., Chang, D., Gonzalez, G.A., King, S.H., Li, D.B., Merchant, A.T., Naghipourfar, M., Nguyen, E., Ricci-Tam, C., Romero, D.W., Sun, G., Taghibakshi, A., Vorontsov, A., Yang, B., Deng, M., Gorton, L., Nguyen, N., Wang, N.K., Adams, E., Baccus, S.A., Dillmann, S., Ermon, S., Guo, D., Ilango, R., Janik, K., Lu, A.X., Mehta, R., Mofrad, M.R.K., Ng, M.Y., Pannu, J., Ré, C., Schmok, J.C., John, J.S., Sullivan, J., Zhu, K., Zynda, G., Balsam, D., Collison, P., Costa, A.B., Hernandez-Boussard, T., Ho, E., Liu, M.-Y., McGrath, T., Powell, K., Burke, D.P., Goodarzi, H., Hsu, P.D., Hie, B.L.: Genome modeling and design across all domains of life with Evo 2. bioRxiv. Pages: 2025.02.18.638918 Section: New Results (2025). 10.1101/2025.02.18.638918. https://www.biorxiv.org/content/10.1101/2025.02.18.638918v1 Accessed 2025-08-17

[22] Hu, W., Ohue, M.: Spatialppiv2: Enhancing protein–protein interaction prediction through graph neural networks with protein language models. Computational and Structural Biotechnology Journal 27, 508–518 (2025) 10.1016/j.csbj.2025.01.022. Accessed 2025-07-19

[23] Liu, D., Young, F., Lamb, K.D., Claudio Quiros, A., Pancheva, A., Miller, C.J., Macdonald, C., Robertson, D.L., Yuan, K.: Plm-interact: extending protein language models to predict protein-protein interactions. Nature Communications 16(1), 9012 (2025) 10.1038/s41467-025-64512-w. Publisher: Nature Publishing Group. Accessed 2025-10-29

[24] Deng, A., Householder, K., Wu, F., Thrun, S., Garcia, K.C., Trippe, B.: Predicting mutational effects on protein binding from folding energy. arXiv. arXiv:2507.05502 [q-bio] (2025). 10.48550/arXiv.2507.05502. http://arxiv.org/abs/2507.05502 Accessed 2025-07-29

[25] Wee, J., Wei, G.-W.: Benchmarking alphafold3’s protein-protein complex accuracy and machine learning prediction reliability for binding free energy changes upon mutation. ArXiv, 2406–039791 (2024). Accessed 2025-07-21

[26] Borkakoti, N., Thornton, J.M.: Alphafold2 protein structure prediction: Implications for drug discovery. Current Opinion in Structural Biology 78, 102526 (2023) 10.1016/j.sbi.2022.102526

[27] Fadini, A., Li, M., McCoy, A.J., Terwilliger, T.C., Read, R.J., Hekstra, D., AlQuraishi, M.: Alphafold as a prior: Experimental structure determination conditioned on a pretrained neural network. bioRxiv (2025) 10.1101/2025.02.18.638828 https://www.biorxiv.org/content/early/2025/03/11/2025.02.18.638828.full.pdf

[28] Mirny, L.A., Shakhnovich, E.I.: Protein structure prediction by threading. why it works and why it does not11edited by f. cohen. Journal of Molecular Biology 283(2), 507–526 (1998) 10.1006/jmbi.1998.2092

[29] Feldman, J., Feldman, T.: Resilient Biosecurity in the Era of AI-Enabled Bioweapons. arXiv. arXiv:2509.02610 [q-bio] (2025). 10.48550/arXiv.2509.02610. http://arxiv.org/abs/2509.02610 Accessed 2025-10-07

[30] Dieckhaus, H., Kuhlman, B.: Protein stability models fail to capture epistatic interactions of double point mutations. bioRxiv, 2024–0820608844 (2024) 10.1101/2024.08.20.608844. Accessed 2025-05-09

[31] Rose, T., Zhou, C., Monti, N.: Affinitylm: Binding-site informed multitask language model for drug-target affinity prediction. In: 2024 IEEE International Conference on Bioinformatics and Biomedicine (BIBM), pp. 727–734 (2024). 10.1109/BIBM62325.2024.10822722. ISSN: 2156-1133. https://ieeexplore.ieee.org/abstract/document/10822722 Accessed 2025-09-03

[32] Gut, J.A., Lemmin, T.: Dissecting alphafold’s capabilities with limited sequence information. bioRxiv (2024) 10.1101/2024.03.14.585076 https://www.biorxiv.org/content/early/2024/06/25/2024.03.14.585076.full.pdf

[33] Agarwal, V., McShan, A.C.: The power and pitfalls of alphafold2 for structure prediction beyond rigid globular proteins. Nature Chemical Biology (2024) 10.1038/s41589-024-01638-w

[34] Li, M.Q.C., Wang, S., Lin, S.-R., Ting, L.E.N., Wan, Z.-H., Xie, G., Zhang, J.: Advantages and limitations of alphafold in structural biology: Insights from recent studies. The Protein Journal (2025) 10.1007/s10930-025-10310-8

[35] Chakravarty, D., Schafer, J.W., Chen, E.A., Thole, J.F., Ronish, L.A., Lee, M., Porter, L.L.: Alphafold predictions of fold-switched conformations are driven by structure memorization. Nature Communications 15(1) (2024) 10.1038/s41467-024-51801-z

[36] Chakravarty, D., Lee, M., Porter, L.L.: Proteins with alternative folds reveal blind spots in alphafold-based protein structure prediction. Current Opinion in Structural Biology 90, 102973 (2025) 10.1016/j.sbi.2024.102973

[37] Molodenskiy, D., Maurer, V.J., Yu, D., Chojnowski, G., Bienert, S., Tauriello, G., Gilep, K., Schwede, T., Kosinski, J.: Alphapulldown2–a general pipeline for high-throughput structural modeling. Bioinformatics 41(3), 115 (2025) 10.1093/bioinformatics/btaf115. Accessed 2025-07-16

[38] Garcia, M., Dixit, S.M., Rocklin, G.J.: Evaluating zero-shot prediction of protein design success by alphafold, esmfold, and proteinmpnn. bioRxiv (2025) 10.1101/2025.07.29.667290 https://www.biorxiv.org/content/early/2025/08/09/2025.07.29.667290.full.pdf

[39] Lin, Z., Akin, H., Rao, R., Hie, B., Zhu, Z., Lu, W., Smetanin, N., Verkuil, R., Kabeli, O., Shmueli, Y., Santos Costa, A., Fazel-Zarandi, M., Sercu, T., Candido, S., Rives, A.: Evolutionary-scale prediction of atomic-level protein structure with a language model. Science 379(6637), 1123–1130 (2023) 10.1126/science.ade2574 https://www.science.org/doi/pdf/10.1126/science.ade2574

[40] Yu, G., Zhao, Q., Bi, X., Wang, J.: Ddaffinity: predicting the changes in binding affinity of multiple point mutations using protein 3d structure. Bioinformatics 40(Supplement 1), 418–427 (2024) 10.1093/bioinformatics/btae232. Accessed 2025-06-08

[41] Feldman, J., Maechler, A., Wang, D., Shakhnovich, E.: Biophysically grounded deep learning improves protein–protein g prediction. bioRxiv (2025) 10.64898/2025.12.23.696257 https://www.biorxiv.org/content/early/2025/12/25/2025.12.23.696257.full.pdf

[42] Zhang, Y., Skolnick, J.: Tm-align: a protein structure alignment algorithm based on the tm-score. Nucleic Acids Research 33(7), 2302–2309 (2005) 10.1093/nar/gki524. Accessed 2024-09-01

[43] Zhang, Y., Skolnick, J.: Scoring function for automated assessment of protein structure template quality. Proteins: Structure, Function, and Bioinformatics 57(4), 702–710 (2004) 10.1002/prot.20264. eprint: https://onlinelibrary.wiley.com/doi/pdf/10.1002/prot.20264. Accessed 2024-12-02

[44] Basu, S., Wallner, B.: Dockq: A quality measure for protein-protein docking models. PLOS ONE 11(8), 0161879 (2016) 10.1371/journal.pone.0161879. Publisher: Public Library of Science. Accessed 2024-11-21

[45] Porter, L.L., Looger, L.L.: Extant fold-switching proteins are widespread. Proceedings of the National Academy of Sciences 115(23), 5968–5973 (2018) 10.1073/pnas.1800168115. Publisher: Proceedings of the National Academy of Sciences. Accessed 2025-12-02

[46] Shah, P., McCandlish, D.M., Plotkin, J.B.: Contingency and entrenchment in protein evolution under purifying selection. Proceedings of the National Academy of Sciences 112(25) (2015) 10.1073/pnas.1412933112

[47] Huss, P., Meger, A., Leander, M., Nishikawa, K., Raman, S.: Mapping the functional landscape of the receptor binding domain of t7 bacteriophage by deep mutational scanning. eLife 10, 63775 (2021) 10.7554/eLife.63775. Publisher: eLife Sciences Publications, Ltd. Accessed 2025-06-26

[48] Woods, H., Schiano, D.L., Aguirre, J.I., Ledwitch, K.V., McDonald, E.F., Voehler, M., Meiler, J., Schoeder, C.T.: Computational modeling and prediction of deletion mutants. Structure 31(6), 713–7233 (2023) 10.1016/j.str.2023.04.005

[49] Guan, C., Wan, F., Torres, M.D.T., Fuente-Nunez, C.d.l.: Improving functional protein generation via foundation model-derived latent space likelihood optimization. bioRxiv. Pages: 2025.01.07.631724 Section: New Results (2025). 10.1101/2025.01.07.631724. https://www.biorxiv.org/content/10.1101/2025.01.07.631724v1 Accessed 2025-05-09

[50] Kempen, M., Kim, S.S., Tumescheit, C., Mirdita, M., Lee, J., Gilchrist, C.L.M., Söding, J., Steinegger, M.: Fast and accurate protein structure search with FoldSeek. Nature Biotechnology (2023)

[51] Lin, Z., Akin, H., Rao, R., Hie, B., Zhu, Z., Lu, W., Santos Costa, A.d., Fazel-Zarandi, M., Sercu, T., Candido, S., Rives, A.: Language models of protein sequences at the scale of evolution enable accurate structure prediction. bioRxiv (2022) 10.1101/2022.07.20.500902 https://www.biorxiv.org/content/early/2022/07/21/2022.07.20.500902.full.pdf

[52] Pak, M.A., Markhieva, K.A., Novikova, M.S., Petrov, D.S., Vorobyev, I.S., Maksimova, E.S., Kondrashov, F.A., Ivankov, D.N.: Using alphafold to predict the impact of single mutations on protein stability and function. PLOS ONE 18(3), 0282689 (2023) 10.1371/journal.pone.0282689. Publisher: Public Library of Science. Accessed 2025-06-22

[53] Ueki, T., Ohue, M.: Antibody complementarity-determining region design using alphafold2 and ddg predictor. The Journal of Supercomputing 80(9), 11989–12002 (2024) 10.1007/s11227-023-05887-9. Accessed 2025-06-22

[54] Wee, J., Wei, G.-W.: Evaluation of alphafold 3’s protein–protein complexes for predicting binding free energy changes upon mutation. Journal of Chemical Information and Modeling 64(16), 6676–6683 (2024) 10.1021/acs.jcim.4c00976. Publisher: American Chemical Society. Accessed 2025-06-22

[55] RCSB Protein Data Bank: biological macromolecular structures enabling research and education in fundamental biology, biomedicine, biotechnology and energy | Nucleic Acids Research| Oxford Academic. https://academic.oup.com/nar/article/47/D1/D464/5144139 Accessed 2024-12-03

[56] Wolf, T., Debut, L., Sanh, V., Chaumond, J., Delangue, C., Moi, A., Cistac, P., Rault, T., Louf, R., Funtowicz, M., Davison, J., Shleifer, S., Platen, P., Ma, C., Jernite, Y., Plu, J., Xu, C., Scao, T.L., Gugger, S., Drame, M., Lhoest, Q., Rush, A.M.: HuggingFace’s Transformers: State-of-the-art Natural Language Processing (2020). https://arxiv.org/abs/1910.03771

[57] Jones, J., Jiang, W., Synovic, N., Thiruvathukal, G.K., Davis, J.C.: What do we know about Hugging Face? A systematic literature review and quantitative validation of qualitative claims (2024). https://arxiv.org/abs/2406.08205

[58] Biasini, M., Schmidt, T., Bienert, S., Mariani, V., Studer, G., Haas, J., Johner, N., Schenk, A.D., Philippsen, A., Schwede, T.: Openstructure: an integrated software framework for computational structural biology. Acta Crystallographica Section D: Biological Crystallography 69(5), 701–709 (2013) 10.1107/S0907444913007051. Publisher: International Union of Crystallography. Accessed 2024-12-03

[59] Mirabello, C., Wallner, B.: Dockq v2: improved automatic quality measure for protein multimers, nucleic acids, and small molecules. Bioinformatics 40(10), 586 (2024) 10.1093/bioinformatics/btae586. Accessed 2024-12-02

