## Supplementary Information for "Adversarial Sequence Mutations in AlphaFold and ESMFold Reveal Nonphysical Structural Invariance, Confidence Failures, and Concerns for Protein Design"

#### S1 Fold-switching protein dataset

For the fifteen experimentally validated fold-switching proteins, mutations were applied with a modified position-weighting scheme that prioritized residues or regions identified in previous studies as inducing conformational transitions. Supplementary Table S2 specifies the mutation ranges and discrete positions for each protein. The weighting formula combined the center bias used for the main 200-protein dataset with an additional multiplier for experimentally validated regions:

$$w_i = \left[\epsilon + (1-\epsilon)\left(1 - \frac{|i-(L-1)/2|}{(L-1)/2}\right)\right] \times r_i$$

where $\epsilon = 0.1$ defines the edge penalty, and $r_i$ is a range multiplier set to 1.5 for positions within specified mutation ranges and 1.0 otherwise. Discrete positions identified in the literature as critical for fold-switching were treated as mandatory mutations and selected first before random sampling. This two-phase approach ensured that experimentally validated switch-inducing residues were always mutated while residues in broader functional regions received a 50% increased probability of selection, all while maintaining the center bias that preferentially targets core positions.

| Metric | Correlation | Coefficient | 5% | 10% | 20% | 40% | 70% | Mean |
|---|---|---|---|---|---|---|---|---|
| TM-score | Pearson | −0.67 | 0.74 | 0.69 | 0.54 | 0.34 | 0.27 | 0.52 |
| | Spearman | −0.73 | — | — | — | — | — | — |

**Table S1 Relationship between mutation percentage and structural prediction accuracy for fold-switching proteins**. TM-score correlation with mutation level and mean metric values at each mutation threshold for ESM3 predictions on a dataset of fifteen experimentally validated fold-switching proteins from Porter et al. [1]. Correlation coefficients indicate the degree of metric degradation as mutation percentage increases.

| PDB (Chain A) | Predicted in [1] | Manual Search | Citation(s) |
|---|---|---|---|
| 2KXO | 1–89 | 2–30, 24, 25 | [2, 3] |
| 2LSH | 29–115 | — | [4] |
| 4OV8 | 247–318 | 128–147, 298–313 | [5] |
| 2MZ7 | 267–312 | 275–280, 306–311 | [6] |
| 4PMK | 27–62 | — | [7] |
| 2N4O | 16–69 | — | [8] |
| 2KTM | 167–201 | 182–217 | [9] |
| 2LE3 | N/A | 19–30, 24 | [10, 11] |
| 2X9C | N/A | 69–70, 71–76 | [12] |
| 3J9E | 2–71 | 1–68, 69–354 | [13] |
| 5SUZ | 474–509, 415–509 | 436; 442–447, 499, 460 | [14] |
| 4HLS | 146–222 | 170, 174 | [15] |
| 1S5P | 48–107, 98–189, 208–274 | — | [16] |
| 3TKA | 236–313 | — | [17] |
| 3GAX | 48–120 | 43–59 | [18] |

**Table S2** Manually identified residues or regions associated with fold switching proteins as collated by [1]. Entries marked with dashes (—) indicate that no precise fold-switching region was found.

| From | To | From | To | From | To | From | To |
|---|---|---|---|---|---|---|---|
| S | W | N | A | K | G | F | T |
| T | F | Q | G | R | A | M | Q |
| C | W | Y | G | W | S | L | N |
| D | W | H | A | A | Q | I | N |
| E | F | G | K | V | E | P | R |

**Table S3** Point mutation transformation pairs used for maximal disruption to protein structures. Standard abbreviations are used for each of the 20 canonical amino acids.

| | Metric | Pearson $r$ | $p$-value | Spearman $\rho$ | $p$-value |
|---|---|---|---|---|---|
| *Monomers* | Top 1 mean | 0.320 | $1.5 \times 10^{-3}$ | 0.352 | $4.4 \times 10^{-4}$ |
| | Top 5 mean | 0.324 | $1.3 \times 10^{-3}$ | 0.333 | $9.2 \times 10^{-4}$ |
| | Top 10 mean | 0.296 | $3.5 \times 10^{-3}$ | 0.308 | $2.3 \times 10^{-3}$ |
| | Top 20 mean | 0.253 | 0.013 | 0.269 | $8.0 \times 10^{-3}$ |
| | Top 100 mean | 0.259 | 0.011 | 0.317 | $1.6 \times 10^{-3}$ |
| | Drop (1→100) | 0.037 | 0.722 | −0.038 | 0.715 |
| *Multimers* | Top 1 mean | 0.268 | $7.7 \times 10^{-3}$ | 0.296 | $3.1 \times 10^{-3}$ |
| | Top 5 mean | 0.278 | $5.7 \times 10^{-3}$ | 0.360 | $2.7 \times 10^{-4}$ |
| | Top 10 mean | 0.294 | $3.3 \times 10^{-3}$ | 0.395 | $5.8 \times 10^{-5}$ |
| | Top 20 mean | 0.333 | $8.1 \times 10^{-4}$ | 0.430 | $9.9 \times 10^{-6}$ |
| | Top 100 mean | 0.382 | $1.1 \times 10^{-4}$ | 0.457 | $2.3 \times 10^{-6}$ |
| | Drop (1→100) | −0.168 | 0.099 | −0.153 | 0.133 |

**Table S4 Correlation between AlphaFold 3 confidence and template neighborhood statistics for monomers ($n = 96$) and multimers ($n = 98$).** Top $N$ refers to the mean query-normalized TM-score of the $N$ best pre-cutoff structural matches. Monomer and multimer confidence is measured by by AlphaFold 3 ranking score.

### References


[1] Porter, L.L., Looger, L.L.: Extant fold-switching proteins are widespread. Proceedings of the National Academy of Sciences **115**(23), 5968–5973 (2018) https://doi.org/10.1073/pnas.1800168115 . Publisher: Proceedings of the National Academy of Sciences. Accessed 2025-12-02

[2] Park, K.-T., Wu, W., Battaile, K.P., Lovell, S., Holyoak, T., Lutkenhaus, J.: The min oscillator uses mind-dependent conformational changes in mine to spatially regulate cytokinesis. Cell **146**(3), 396–407 (2011)

[3] Ayed, S.H., Cloutier, A.D., McLeod, L.J., Foo, A.C., Damry, A.M., Goto, N.K.: Dissecting the role of conformational change and membrane binding by the bacterial cell division regulator mine in the stimulation of mind atpase activity. Journal of Biological Chemistry **292**(50), 20732–20743 (2017)

[4] Morris, V.K., Kwan, A.H., Sunde, M.: Analysis of the structure and conformational states of dewa gives insight into the assembly of the fungal hydrophobins. Journal of molecular biology **425**(2), 244–256 (2013)

[5] Lukoyanova, N., Kondos, S.C., Farabella, I., Law, R.H., Reboul, C.F., Caradoc-Davies, T.T., Spicer, B.A., Kleifeld, O., Traore, D.A., Ekkel, S.M., *et al.*: Conformational changes during pore formation by the perforin-related protein pleurotolysin. PLoS biology **13**(2), 1002049 (2015)

[6] Kadavath, H., Jaremko, M., Jaremko, Ł., Biernat, J., Mandelkow, E., Zweckstetter, M.: Folding of the tau protein on microtubules. Angewandte Chemie International Edition **54**(35), 10347–10351 (2015)

[7] Hamiaux, C., Maddumage, R., Middleditch, M.J., Prakash, R., Brummell, D.A., Baker, E.N., Atkinson, R.G.: Crystal structure of kiwellin, a major cell-wall protein from kiwifruit. Journal of structural biology **187**(3), 276–281 (2014)

[8] Pham, C.L., Rey, A., Lo, V., Soulès, M., Ren, Q., Meisl, G., Knowles, T.P., Kwan, A.H., Sunde, M.: Self-assembly of mpg1, a hydrophobin protein from the rice blast fungus that forms functional amyloid coatings, occurs by a surface-driven mechanism. Scientific reports **6**(1), 25288 (2016)

[9] Adrover, M., Pauwels, K., Prigent, S., Chiara, C., Xu, Z., Chapuis, C., Pastore, A., Rezaei, H.: Prion fibrillization is mediated by a native structural element that comprises helices h2 and h3. Journal of Biological Chemistry **285**(27), 21004–21012 (2010)

[10] Rao, J.N., Warren, G.Z., Estolt-Povedano, S., Zammit, V.A., Ulmer, T.S.: An environment-dependent structural switch underlies the regulation of carnitine palmitoyltransferase 1a*. Journal of Biological Chemistry **286**(49), 42545–42554 (2011)

[11] Rao, J.N., Warren, G.Z., Estolt-Povedano, S., Zammit, V.A., Ulmer, T.S.: An environment-dependent structural switch underlies the regulation of carnitine palmitoyltransferase 1a*. Journal of Biological Chemistry **286**(49), 42545–42554 (2011)

[12] Poyraz, Ö., Schmidt, H., Seidel, K., Delissen, F., Ader, C., Tenenboim, H., Goosmann, C., Laube, B., Thünemann, A.F., Zychlinsky, A., *et al.*: Protein refolding is required for assembly of the type three secretion needle. Nature structural & molecular biology **17**(7), 788–792 (2010)

[13] Zhang, X., Patel, A., Celma, C.C., Yu, X., Roy, P., Zhou, Z.H.: Atomic model of a nonenveloped virus reveals ph sensors for a coordinated process of cell entry. Nature structural & molecular biology **23**(1), 74–80 (2016)

[14] Gammons, M.V., Renko, M., Johnson, C.M., Rutherford, T.J., Bienz, M.: Wnt signalosome assembly by dep domain swapping of dishevelled. Molecular cell **64**(1), 92–104 (2016)

[15] Sweeting, B., Brown, E., Khan, M.Q., Chakrabartty, A., Pai, E.F.: N-terminal helix-cap in $\alpha$-helix 2 modulates $\beta$-state misfolding in rabbit and hamster prion proteins. PloS one **8**(5), 63047 (2013)

[16] Zhao, K., Chai, X., Marmorstein, R.: Structure and substrate binding properties of cobb, a sir2 homolog protein deacetylase from escherichia coli. Journal of molecular biology **337**(3), 731–741 (2004)

[17] Wei, Y., Zhang, H., Gao, Z.-Q., Wang, W.-J., Shtykova, E.V., Xu, J.-H., Liu, Q.-S., Dong, Y.-H.: Crystal and solution structures of methyltransferase rsmh provide basis for methylation of c1402 in 16s rrna. Journal of structural biology **179**(1), 29–40 (2012)

[18] Szymańska, A., Jankowska, E., Orlikowska, M., Behrendt, I., Czaplewska, P., Rodziewicz-Motowidło, S.: Influence of point mutations on the stability, dimerization, and oligomerization of human cystatin c and its l68q variant. Frontiers in Molecular Neuroscience **5**, 82 (2012)
